# Homogeneous environmental selection overturns distance-decay relationship of soil prokaryotic community

**DOI:** 10.1101/2021.05.13.443991

**Authors:** Biao Zhang, Kai Xue, Shutong Zhou, Kui Wang, Wenjing Liu, Cong Xu, Lizhen Cui, Linfeng Li, Qinwei Ran, Ronghai Hu, Yanbin Hao, Xiaoyong Cui, Yanfen Wang

## Abstract

Though being fundamental to global diversity distribution, little is known about the geographic pattern of soil microorganisms across different biomes on a large scale. Here, we investigated soil prokaryotic communities from Chinese northern grasslands on a scale up to 4,000 km in both alpine and temperate biomes. Surprisingly, prokaryotic similarities increased with geographic distance after tipping points of 1,760 - 1,920 km, overturning the well-accepted distance-decay relationship and generating a significant U-shape pattern. Such U-shape pattern was likely due to decreased disparities in environmental heterogeneity along with geographic distance when across biomes, as homogeneous environmental selection dominated prokaryotic assembly based on βNTI analysis. Consistently, short-term environmental heterogeneity also followed the U-shape pattern spatially, mainly attributed to dissolved nutrients. In sum, these results demonstrate that homogeneous environmental selection via dissolved nutrients overwhelmed the “distance” effect when across biomes, subverting the previously well-accepted geographic pattern for microbes on a large scale.

## Introduction

To clarify the spatial pattern of biodiversity is one of primary aims in ecology and biogeography (1, 2). In past decades, intensive biogeographic studies have been conducted for macro-organisms, including plants (3–8), insects (9, 10) and animals (11, 12). With the emergence and development of next-generation sequencing, increasing attention has been paid recently to the spatial pattern of microorganisms. The similarity of microbial communities has been observed to decrease as that of macro-organisms does with geographic distance, so-called distance-decay relationship, in different habits (e.g., forests (13, 14), grasslands (15), deserts (16) and agriculture soils (17–19)) for bacteria (15, 20, 21), archaea (18, 22), fungi (23–26) and speicific micorbial functional groups (e.g. ammonia-oxidizing archaea, ammonia-oxidizing and sulfate-reducing bacteria (27–30)). The reported distance-decay relationship has been regarded as a principal generalization in nature, which generally rejects the hypothesis of “everything is everywhere, but environment selects” (31).

However, no consensus has been reached so far on underlying mechanisms of the distance-decay relationship for soil microbial communities. A few mechanisms for biodiversity maintenance (20) were proposed to be responsible for such relationship as well, including dispersal limitation, environmental heterogeneity and stochastic processes (4, 32, 33). Microorganisms have been observed to have the dispersal limitation as macro-organisms do (2, 34, 35), crucial in biodiversity maintenance and evolution (33, 36). Spatial configurations and the nature of landscapes influence the dispersal rate of organisms among sites (32), and communities tend to be more similar in open and topographically homogeneous settings than in heterogeneous landscapes. Moreover, environmental heterogeneity tends to increase with geographic distance, responsible for the distance-decay relationship as well (2, 18). Communities are expected to become increasingly different along with geographic distance as their species are sorted according to their niche requirements (34). Under this scenario, dissimilarities among communities parallel to increasing disparities in environmental heterogeneity along with geographic distance. Furthermore, stochastic processes in birth, death, migration, disperse and drift may also contribute to the distance-decay relationship for soil microbial communities (37–39).

The relative importance of various mechanisms is still unclear (4, 32, 33), likely being scale- and biome-dependent. Environment heterogeneity has been reported to be more important in influencing the spatial distribution of microorganisms at local scales up to hundreds of kilometers (19, 40, 41), while dispersal limitation dominated the distance-decay relationship on larger scales (42, 43). Moreover, different biomes under distinct climate conditions/latitudes may have different turnover rates for community similarity over geographic distance. High temperature in forest soils was reported to lead to a lower turnover rate (30), likely due to accelerated biochemical reactions and increased ecological niche breadth (44). Community similarity was observed to decline faster at high than low latitudes on large scales, while the turnover rate was higher at low latitudes on small scales (32). However, most previous studies were conducted locally or regionally within a single biome or climate type. Thus, surveys for the geographic pattern of soil microorganisms across different biomes is essential to understand the spatial pattern of microbial communities beyond these scales.

Here, we collected grassland soil samples from two biomes with distinct hydrothermal conditions to investigate the spatial pattern of prokaryotic communities and underlying mechanisms. A total of 258 samples were collected from the top- and subsoils in alpine and temperate biomes on a scale up to 4,000 km, on Tibet Plateau and Inner Mongolia Plateau, respectively. Our objectives were to test the following hypotheses: (I) soil prokaryotic community similarity would decrease over geographic distance within and across biomes; (II) the turnover rate of soil prokaryotic community similarity over geographic distance would be higher in the temperate than alpine biome, as temperate biome is with a wider temperature range; (III) the turnover rate of soil prokaryotic community similarity over geographic distance would be lower in top- than subsoil, as the subsoil may be less dynamic and affected by environmental factors like UV and wind.

## Results

### Prokaryotic and plant community similarity over geographic distance

A total of 11,063 OTUs were detected from all 258 grassland soil samples in both alpine and temperate biomes. A significant (*P* < 0.001) binomial relationship (U shape) was observed for the prokaryotic community over geographic distance in top- (R^2^ = 0.161) or subsoil (R^2^ = 0.114) from all sites on a scale up to 4,000 km. Specifically, the prokaryotic community similarity in topsoil decreased over geographic distance on a scale of < 1,920 km mostly within either temperate or alpine biome, but increased after this tipping point when across biomes (in pairwise sites between alpine and temperate biomes). Similarly, the prokaryotic community similarity in subsoil decreased over geographic distance on a scale of < 1,760 km mostly within either temperate or alpine biome, but increased after this tipping point when across biomes (Figure 1). When across biomes, the prokaryotic community similarity increased significantly over geographic distance with similar slopes (turnover rates) in top- (slope = 0.007, R^2^ = 0.199) and subsoil (slope = 0.006, R^2^ = 0.134).

**Figure 1.**
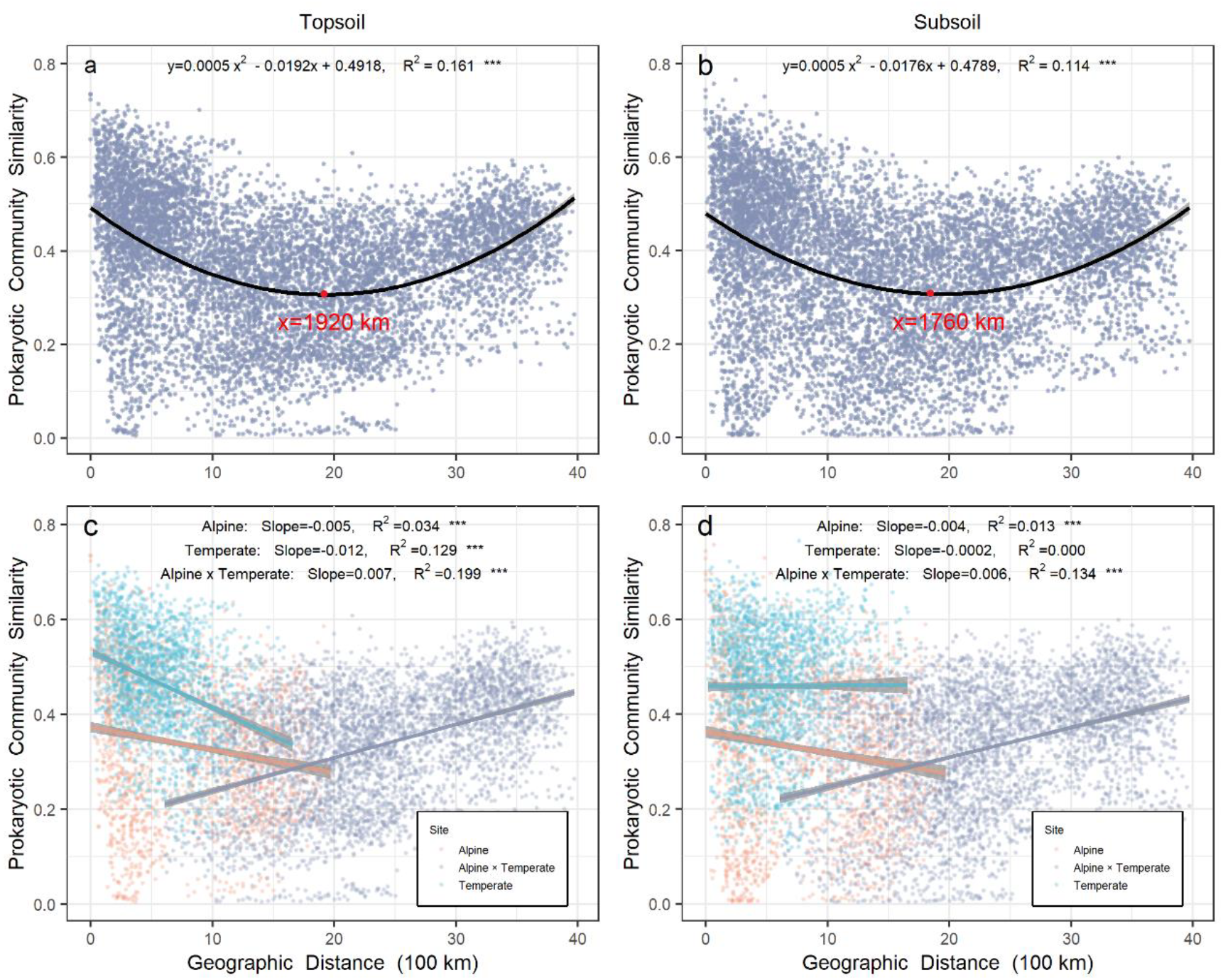
Relationship between prokaryotic community similarity over geographic distance in Chinese northern grassland. Panel **a** and **c** represent the prokaryotic community in topsoil, while panel **b** and **d** represent the prokaryotic community in subsoil. Orange and light blue points represent pairwise sites within the alpine and temperate biome, respectively. Grey points represent pairwise sites between the alpine biome cross temperate biome. Grey shades stand for 95% confidence interval.

Within the alpine biome, a valid (*P* < 0.001) distance-decay relationship was observed for the prokaryotic community over geographic distance in top- (R^2^ = 0.034) or subsoil (R^2^ = 0.013). However, within the temperate biome, the distance-decay relationship for the prokaryotic community occurred only in topsoil (R^2^ = 0.129, *P* < 0.001), while no relationship was observed in subsoil. In topsoil, prokaryotic community similarity had a higher turnover rate in the temperate (− 0.012) than alpine biome (− 0.005).

Similar to the prokaryotic community, the plant community also exhibited a significant U-shape relationship (R^2^ = 0.071, *P* < 0.001) for its similarity over geographic distance in all sites on a scale up to 4,000 km, with a tipping point of 1,858 km (Figure S2a). A significant (*P* < 0.001) distance-decay relationship for plant community was observed within the alpine (Figure S2b, R^2^ = 0.015, *P* < 0.001) or temperate biome (Figure S2b, R^2^ = 0.005, *P* < 0.01).

### Prokaryotic community similarity over environmental distance

Relatively short-term environmental similarity also exhibited a U-shape pattern over geographic distance in either top- or subsoil from all sites on a scale up to 4,000 km (Figure 2 a and d). In contrast, relatively long-term environmental similarity, much higher than short-term environmental similarity on the same scale, did not change greatly over geographical distance in either top- or subsoil.

**Figure 2.**
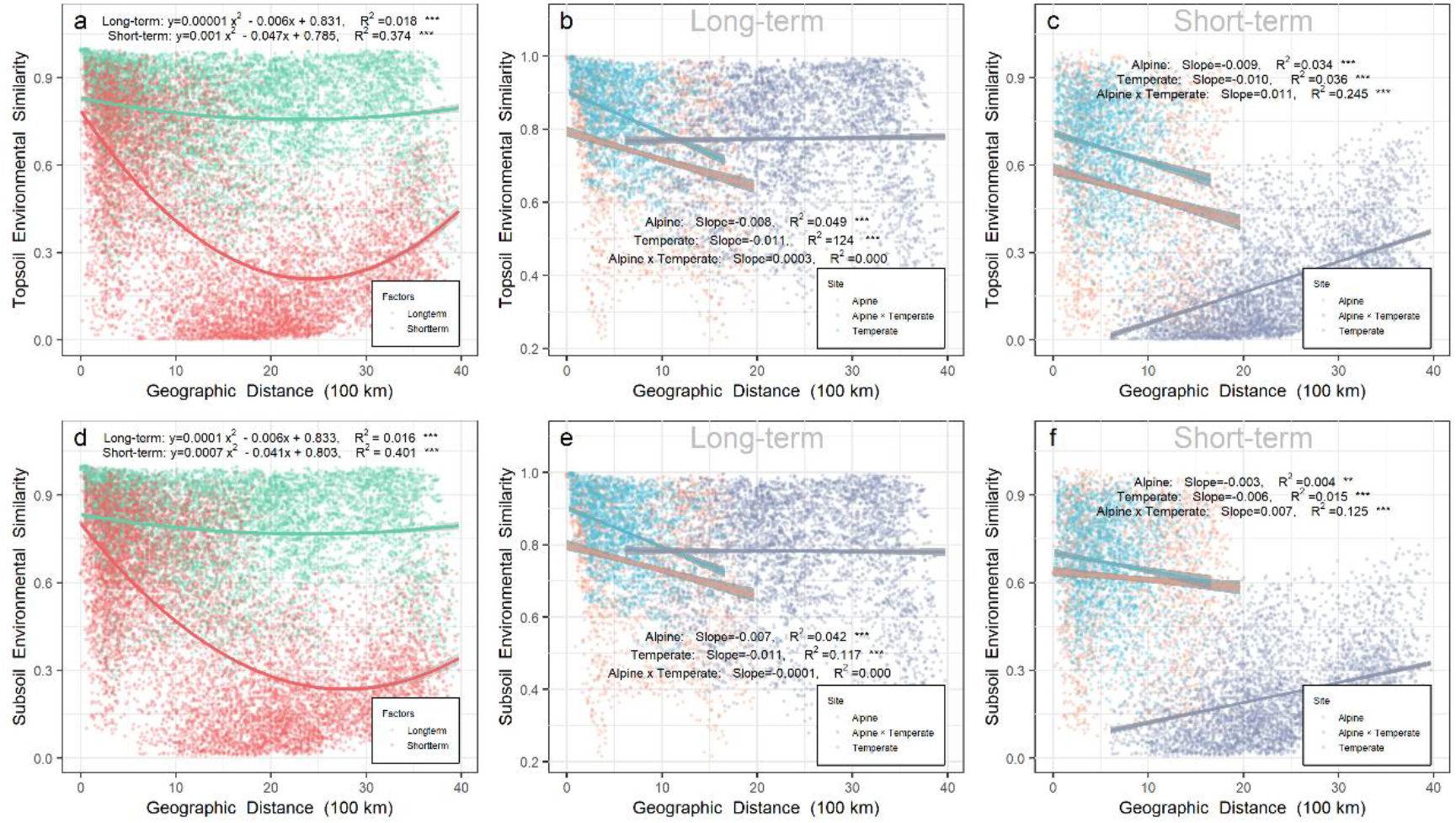
Relationship between environmental similarity over geographic distance in Chinese northern grassland. Panel a, b and c represent the environmental similarity in topsoil, while panel d, e and f represent the environmental similarity in subsoil. The environmental similarity was calculated by Bray-Curtis index based on relatively long-term (b, e & green points in a and d) or short-term variables (c, f & red points in a and d). Relatively long-term environmental variables included mean annual precipitation (MAP), mean annual temperature (MAT), pH, soil organic carbon (SOC), soil total nitrogen (TN) and soil total phosphorus (TP). Relatively short-term environmental variables included soil water content (SWC), soil available phosphorus (AP), dissolved organic carbon (DOC), dissolved organic nitrogen (DON), NH_4_^+^, and NO_3_^−^. Orange, grey, and light blue points represent pairwise sites within the alpine biome, within temperate biome, and alpine biome cross temperate biome, respectively. Grey shades stand for 95% confidence interval.

Soil prokaryotic community similarity decreased significantly (*P* < 0.001) in all sites over the relatively long-term (turnover rate = − 0.291 or − 0.278 in top- or subsoil, respectively) or short-term (turnover rate = − 0.193 or − 0.159 in top- or subsoil, respectively) environmental distance (Figure S3 a and e). In the topsoil, the prokaryotic community similarity decreased significantly (*P* < 0.001) over the relatively long-term (turnover rate = − 0.277, R^2^ = 0.131) or short-term (turnover rate = − 0.194, R^2^ = 0.135) environmental distance within the alpine biome (Figure S3 b), as well as decreased over long-term (turnover rate=−0.339, R^2^ = 0.108)or short-term(turnover rate=−0.063, R^2^ = 0.011)environmental distance within the temperate biome (Figure S3 c). In the subsoil, the prokaryotic community similarity decreased significantly over the relatively long-term (turnover rate = − 0.279, R^2^ = 0.104) or short-term (turnover rate = − 0.215, R^2^ = 0.093) environmental distance within the alpine biome (Figure S3 f), while there was no relationship within the temperate biome (Figure S3 g). In pairwise sites between the alpine cross temperate biome (Figure S3 d and h), the prokaryotic community similarity decreased over relatively long-term (turnover rate = − 0.161 or − 0.130 in top- or subsoil, respectively) or short-term (turnover rate = − 0.191 or − 0.175 in top- or subsoil, respectively) environmental distance.

As revealed by Partial Mantel test (Table 1), the significant decay relationship between topsoil prokaryotic community similarity and relatively short-term environmental distance across biomes was mainly driven by soil water content (SWC, r = 0.167, p = 0.002), available phosphorus (AP, r = 0.163, p = 0.004), dissolved organic carbon (DOC, r = 0.221, p < 0.001) and dissolved organic nitrogen (DON, r = 0.225, p < 0.001). Similar short-term environmental variables (except AP) were responsible for the significant decay relationship in the subsoil.

**Table 1.** Partial Mantel test for relationship between prokaryotic community similarity and relatively short-term environmental variables across biomes.

|  | Alpine × Temperate |  |  |  |
| --- | --- | --- | --- | --- |
|  | Topsoil |  | Subsoil |  |
|  | r | p | r | p |
| SWC | 0.167 | **0.002** | 0.225 | **<0.001** |
| AP | 0.163 | **<0.001** | 0.038 | 0.231 |
| DOC | 0.221 | **<0.001** | 0.117 | **0.027** |
| DON | 0.225 | **<0.001** | 0.200 | **0.002** |
| NH<sub>4</sub><sup>+</sup> | 0.026 | 0.331 | 0.065 | 0.093 |
| NO<sub>3</sub><sup>-</sup> | 0.068 | 0.140 | 0.024 | 0.323 |
| Short-term environment factors* | 0.273 | **0.001** | 0.219 | **0.001** |

\*by Mantel test

Within the alpine biome, the significant distance-decay relationship between topsoil prokaryotic community similarity and relatively long-term environmental distance was mainly driven by mean annual precipitation (MAP, r = 0.249, p < 0.001), soil organic carbon (SOC, r = 0.269, p < 0.001), and soil total nitrogen (TN, r = 0.239, p < 0.001), while SWC (r = 0.534, p < 0.001), AP (r = 0.297, p < 0.001), DOC (r = 0.285, p < 0.001), DON (r = 0.278, p < 0.001), NH_4_^+^ (r = 0.223, p < 0.001), and NO_3_^−^ (r = 0.133, p = 0.031) were responsible for the significant distance-decay relationship between topsoil prokaryotic community similarity and relatively short-term environmental distance (Table S1 and S2, Figure S4). Similar relatively long-term and short-term environmental variables (except AP, NH_4_^+^, and NO_3_^−^ were responsible for the significant distance-decay relationship of prokaryotic community similarity in the subsoil within the alpine biome. Within the temperate biome, the significant distance-decay relationship between topsoil prokaryotic community similarity and relatively long-term environmental distance was driven by MAP (r = 0.334, p < 0.001), SOC (r = 0.128, p = 0.022), and TN (r = 0.117, p = 0.021), while DOC (r = 0.059, p = 0.034) was responsible for the significant distance-decay relationship between topsoil prokaryotic community similarity and relatively short-term environmental distance.

### Deterministic and stochastic processes in prokaryotic community assembly

*β*NTI (*β*-nearest taxon index) analysis was used to distinguish different processes in prokaryotic community assembly. As shown in Figure 3, the range of |*β*NTI| > 2 indicated that deterministic processes played a dominant role (>85%) in prokaryotic community assembly. The contribution of deterministic processes was relatively lower in the alpine (Figure 3j; 92.91% and 87.47% in top- and sub-soil, respectively) than temperate (96.63% and 94.04% in top- and sub-soil, respectively) biome, and higher in the topsoil than subsoil in all sites. Moreover, most*β*NTI values were less than −2 either in the top- or subsoil from all sites, indicating that prokaryotic communities were assembled mainly by homogeneous selection in deterministic processes.

**Figure 3.**
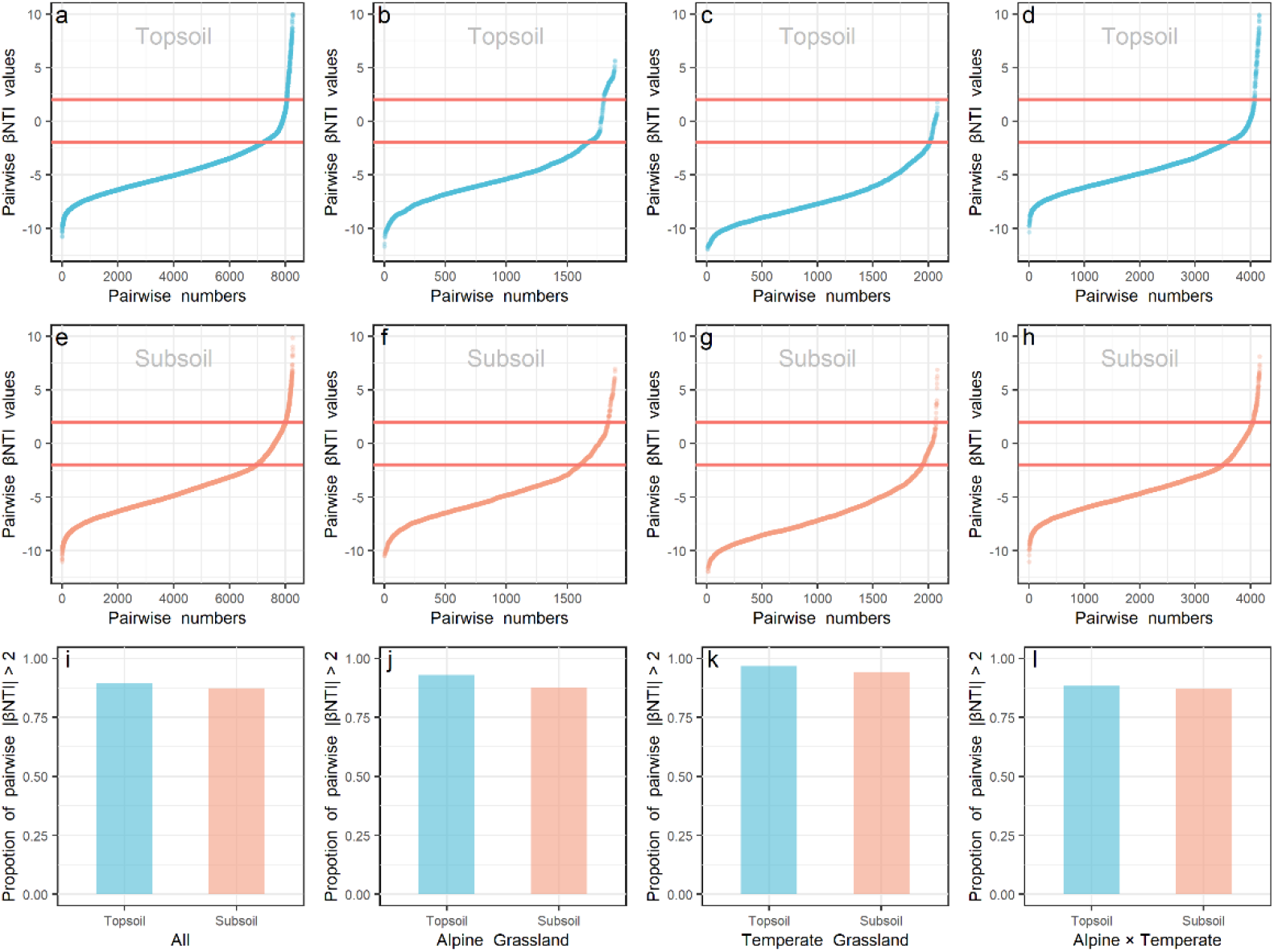
*β*NTI values. *β*NTI values of prokaryote in the top- (a, b, c and d) and subsoil (e, f, g and h) from all sites (a and e), within alpine (b and f) or temperate biome (c and g), and pairwise sites in the alpine cross temperate biomes (d and h) were presented. The proportion of |*β*NTI| >2 (deterministic processes) in the top- and subsoil from all sites (i), within alpine biome (j), within temperate biome (k) and pairwise sites in the alpine cross temperate biomes (l) were presented. Light blue and orange colors stand for βNTI values and their proportions in the top- and subsoil, respectively.

We further compared the immigration rates (*m*) of prokaryotes (Figure S7) based on the algorithm developed by Hubbell for the neutral theory (49). Prokaryotic immigration rates were significantly lower in the alpine (0.159 ± 0.008 and 0.146 ± 0.008 in top+ and subsoil, respectively) than temperate biome (0.261 ± 0.010 and 0.246 ± 0.009 in top+ and subsoil, respectively) in the same soil layer. Moreover, immigration rates were higher in the top+ (0.159 ± 0.008 and 0.261 ± 0.010 in the alpine and temperate biomes, respectively) than subsoil (0.146 ± 0.008 and 0.246 ± 0.009 in the alpine and temperate biomes, respectively) in the same biome.

In the pairwise sites between alpine cross temperate biomes, structural equation model (SEM; Figure 4) showed prokaryotic community similarity was mainly affected by geographic distance (r = 0.388 and 0.320 in top- and subsoil, respectively), relatively long-term environmental distance (r = −0.171 and −0.130 in top- and subsoil, respectively), plant community dissimilarity (r = −0.124 and −0.065 in top- and subsoil, respectively), and relatively short-term environmental distance (r = −0.046 and −0.069 in top- and subsoil, respectively).

**Figure 4.**
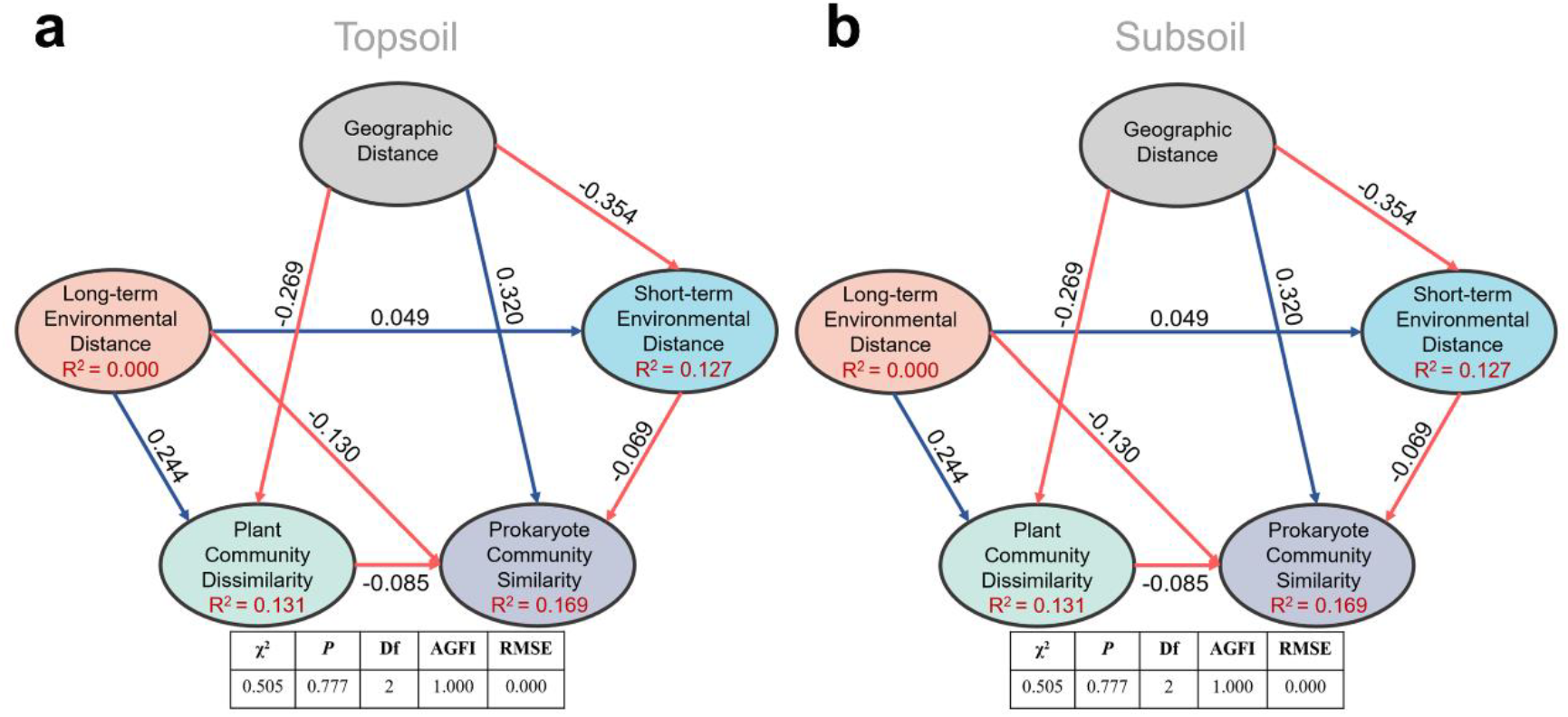
Structure equation model to quantify effects of geographic distance, relatively long225 term and short-term environmental distance, and plant community dissimilarity on soil prokaryotic community similarity in the pairwise sites between alpine cross temperate biomes, either in. top-**(a)** or subsoil **(b)**. Red and blue lines stand for negative and positive correlations, respectively; bold lines stand for significance at p = 0.05 level.

Within each biome, geographic distance only had a direct effect on topsoil prokaryotic community similarity (r = −0.275) in the temperate grassland, while its effect in topsoil of the alpine biome was only indirect through relatively short-term (r = 0.140) and long-term environmental distances (r = 0.222), as well as plant community dissimilarity (r = 0.232). In the alpine biome, increases in plant community dissimilarity, relatively long-term and short-term environmental distances directly decreased the similarity of soil prokaryotic community. The explained variances of soil prokaryotic community similarity were 26.8% and 21.3% in the top- and subsoil, respectively (Figure S6). In the temperate biome, other than the direct effect of geographic distance, plant community dissimilarity, relatively long-term and short-term environmental distances also affected topsoil prokaryotic community similarity directly. In the subsoil of the temperate biome, prokaryotic community similarity was not significantly correlated with any factors and the explained variance was merely 0.2%.

## Discussion

“Everything is related to everything else, but near things are more related to each other” is termed as “the first law of geography” (45). Ecologists and biogeographers refer it to the negative relationship between community similarity and distance as a geographical distance-decay relationship (46, 47). Though being regarded as a principal generalization, distance-decay relationship was denied and overturned in this study as prokaryotic community similarity increased with the geographical distance after tipping points of 1,920 - 1,760 km when across biomes. This finding is contradictory with most previous studies, including a report that was conducted even at the similar scale of 4,000 km to ours but within a single biome (temperate biome) (15). Consistently, when within a single biome of alpine or temperate grassland, the distance-decay relationship was also valid in this study.

However, over environmental distance, prokaryotic community similarity showed significant decay relationships (Figure S5) in all sites on a scale up to 4,000 km in this study, consistent with previous reports (48, 49). Thus, the U-shape pattern of prokaryotic community similarity over geographic distance in all sites may be attributed to disparities in environmental heterogeneity over geographic distance. This is also supported by the *β*NTI analysis showing that the prokaryotic community assembly was dominantly determined by homogeneous selection in deterministic processes, referring similar habitats (environment) harbor similar prokaryotic communities (50, 51). The role of environment filtering, including biotic and abiotic factors, on microbial community assembly has been widely reported at the scale of hundreds to thousands of kilometers (19, 40, 41).

Environmental variables measured in this study were separated into relatively long-term and short-term environmental variables, judged by dynamic time. Interestingly, the similarity of relatively short-term environmental variables exhibited a U shape pattern over geographic distance in all sites on a scale up to 4,000 km, while that of relatively long-term environmental variables did not, indicating that relatively short-term environmental variables may be responsible for the homogeneous selection shaping the U shape pattern of prokaryotic community over geographic distance. Consistently, SEM analyses also revealed a significant direct effect of relatively short-term environmental distance on prokaryotic community similarity when across biomes. The Partial Mantel test further demonstrated that water (MAP, SWC) and dissolved nutrients (DOC, DON) were the primary short-term environmental factors in the significant relationship between prokaryotic community similarity and geographic distance. The effects of water on the microbial community have been widely reported (52, 53), as soil water content could determine soil texture, bulk density, oxygen availability and connectivity within soils (54–56), which can vitally influence soil microbial community composition (57) and microbial basal respiration in the semi-arid area (58). Soil water content also influences microbial communities through changing nutrient availability. Plenty of studies found that DOC and DON affected the distribution pattern of soil microbes (41, 59–62). Compared to other nutrients, DOC and DON can be utilized by microbes more directly and easily to provide energy and nutrients for supporting their growth (63, 64).

In addition to abiotic factors, plant community attributes (65–69), especially plant species identity (48, 70, 71), may also be important for the U-shape pattern of prokaryotic community similarity. Our results demonstrated that soil prokaryotic community similarity decreased over the plant community dissimilarity in all sites on a scale up to 4,000 km. Moreover, SEM also revealed the significantly direct effect of plant community dissimilarity on prokaryotic community similarity in pairwise sites between alpine cross temperate biomes. Plant community composition and diversity can affect soil prokaryotic communities through altering the quality and quantity of organic matter input to soils by the forms of litterfall and root exudates (72). Plants exude a substantial proposition (11 - 40%) of photosynthesis-derived carbon (73), including sugars, amino acids, organic acids, fatty acids and secondary metabolites (73–75). Their compounds in exudation can attract beneficial microorganisms deliberately and influence the assembly of soil microbiomes to promote plants’ adaptation to the surrounding environment (76–81).

In the topsoil of temperate biome, MAP rather than MAT was responsible for the significant decay relationship between prokaryotic community similarity and long-term environmental distance, together with SOC and TN. In the alpine biome, both MAP and MAT were responsible for the significant distance-decay relationship between the prokaryotic community similarity and long-term environmental distance in either top- or subsoil, together with pH, SOC, and TN. These phenomena indicated that the spatial pattern of soil prokaryotic community was driven by both temperature and water in the alpine biome with a low temperature range (−0.8 to 5.9 °C) and a low precipitation range (84 to 528 mm) (82). However, the spatial pattern of soil prokaryotic community in the temperate biome was driven by precipitation (159 to 460 mm) only in the topsoil (15), where temperatue is a limting factor (1.23 to 4.4 °C). Consistently, our previous study demonstrated that soil microbial diversity in the alpine biome was mainly affected by temperature especially under the condition of limited precipitation (83).

In the topsoil, the turnover rate of prokaryote was higher in temperate than alpine biome (Figure 1). We found that the immigration rate (*m*) of topsoil prokaryotes was higher in the temperate biome than that of the alpine biome (Figure S7), indicating a weakened dispersal limitation that may be responsible for the higher similarity of prokaryotic community in the temperate biome (2, 50). The effects of dispersal limitation on microbial communities (43, 84, 85) were dependent on ecosystems or environmental habitats (86–88). Harsh environments (50, 51, 89) with low temperature, high UV and complex mountain terrain in the alpine biome on the Tibet Plateau would not be conducive for soil prokaryote to disperse. In contrast, the temperate biome has benign temperature, low UV and better landscape connectivity to promote the spatial dispersal of microorganisms.

Notably, within the temperate biome across 1,661 km, prokaryotic community similarity did not change with the geographic distance in the subsoil. Moreover, subsoil prokaryotic community similarity was not linked with plant community dissimilarity (Figure S5) and long-term environmental distance (Figure S3), or correlated weakly with short-term environmental distance based on the correlation test (R^2^ = 0.011) and SEM (r = − 0.002), denying the possibility of plant dependent and environment heterogeneity driven. Cases of no distance-decay relationship for microorganism have also been reported previously (29, 90, 91), explained by the absence of dispersal limitation (29), paleogeographic history (90, 91) that may also be applied in our case and plant dependent (91).

## Conclusion

This study provides a systematical analysis of the spatial pattern of soil prokaryotic communities in the northern grassland of China. Soil prokaryotic similarity exhibited a U-shape distribution pattern over geographic distance at a scale of up to 4,000 km. This finding overturns the well-accepted distance-decay relationship, which was only valid in the top- or subsoils within the alpine biome and topsoil only within the temperate biome. Despite different climate and ecosystem types in the alpine and temperate biomes, habitats far more apart when across biomes were more similarly as revealed by the U-shape pattern for short-term environmental factors over geographic distance. Consistently, deterministic processes were found to dominate the soil prokaryotic community assembly by *β*NTI analysis, and further partial Mantel analysis revealed that water (MAP and SWC) and dissolved nutrients (DOC and DON) together may be responsible for the U pattern of prokaryotic community similarity over geographic distance, overturning the distance-decay relationship.

## Materials and Methods

### Study sites and field sampling

A total of 129 sites and 258 samples were collected from the northern grassland of China. Among them, 128 samples from 64 sites (red dots in Figure S1) were collected from July 29 to August 14, 2014 in the Qinghai-Tibet Plateau alpine biome, China. Alpine biome sample sites covered a variety of alpine ecosystems, including alpine meadow, alpine steppe, alpine desert, and shrub. One hundred and twenty-eight samples from 65 sites (blue dots in Figure S1) were collected from September 10 to 24, 2015 in the Inner Mongolia temperate biome, China. Temperate biome sample sites covered three types of temperate biome ecosystem, namely temperate meadow, temperate steppe, and temperate desert. The distance between each two adjacent sample sites was no less than 60 km and removed from potential human interference such as towns, villages, and roads.

A GPS (global positioning system) was used to record the geographic coordinates and altitudes of each sample site. Five plots were selected randomly at each site and the distance between the adjacent two plots was no less than 10 m. After removing plant aboveground biomass and litter, three topsoil (0-5 cm in depth) and subsoil (5-20 cm in depth) cores (7 cm in diameter) were randomly sampled within each plot.

Topsoil or subsoil samples from each plot were pooled and then sieved through a 2 mm mesh, and the roots were selected as belowground biomass (BGB). Sieved topsoil or subsoil samples were divided into two subsamples. One part was stored at room temperature and dried in the shade for measuring physical and chemical properties. The other part was stored at approximately 4 °C in the field by a mobile refrigerator, delivered with dry ice to the laboratory in Beijing, and finally frozen at −80 °C in a freezer before DNA extraction.

### Soil properties

Soil pH was measured by pH meter (STARTER3100, Ohaus Instruments Co., Ltd., Shanghai, China) with a 1:5 of soil water ratio (5 g soil: 25 mL ddH_2_O). SWC was measured by ovening fresh soil samples at 105°C for 24 h. SOC was measured by a TOC analyzer (Liqui TOC II; Elementar Analysensysteme GmbH, Hanau, Germany). Soil TN was measured on an auto-analyzer (SEAL Analytical GmbH, Norderstedt, Germany). Soil TP and AP were measured by a UV-VIS spectrophotometer (UV2700, SHIMADZU, Japan). Nitrate-N (NO_3_^−^) and ammonium-N (NH_4_^+^) were extracted with 2 M KCl (soil mass to solution ratio of 1:5) and then analyzed on a continuous-flow ion auto-analyzer (SEAL Analytical GmbH, Norderstedt, Germany). Soil DOC and DON were measured on a TOC Analyser (Liqui TOC II; Elementar Analysensysteme GmbH, Hanau, Germany). Plant aboveground biomass (AGB) and belowground biomass (BGB) were measured after oven drying at 65 °C for 72 h. MAT and MAP of each study site were obtained from “China Meteorological Data Service Center” (CMDC: https://data.cma.cn/) by latitude and longitude.

### Microbial analysis

Soil genomic DNA was extracted from 0.25 g frozen soil three times at each soil layer at each site and then mixed into one DNA sample using PowerSoil™ DNA Isolation Kits (MO BIO Laboratories, Carlsbad, CA, USA). The quality of extracted DNA was assessed based on OD 260/280 nm and 260/230 nm absorbance ratios by NanoDrop (2000) spectrophotometer (NanoDrop Technologies inc., Wilmington, DE, USA). Primer pair 515F (5’-GTGYCAGCMGCCGCGGTA-3’) and 909R (5’-CCCCGYCAATTCMTTTRAGT-3’) was selected to amplify the V4-V5 region of 16S rRNA and the target fragment length was 374 bp, and the 12bp barcode was added at the end of 5’ of 515F. A 50 μL PCR reaction system was configured in 0.2 mL tube, including 2 μL template DNA diluent, 4 μL dNTP, 4 μL Mg_2_^+^, 5 μL Buffer, 0.5 L Ex Taq™ enzyme, 1 μL forward primer, 1 μL reverse primer, 32.5 L ddH_2_O. The PCR procedure was performed as follows: predenaturation at 95°C for 10 min, 30 PCR cycles (deformation at 94°C for 30 s, annealing at 53°C for 25 s, extension at 68°C for 45 s), and a final extension at 72°C for 10 min.

The PCR products were purified by 1% agarose gel using GeneJET Gel Extraction Kit (Thermo, USA). The purified DNA was tested by NanoDrop (2000) spectrophotometer (NanoDrop Technologies inc., Wilmington, DE, USA). All purified DNA samples were mixed in 100 ng before database construction and sequencing, which was performed by Illumina Miseq in Chengdu Biology Institute.

The MiSeq raw data was analyzed by UPARSE pipeline with USEARCH 8 software to obtain an operational taxon units (OTU) table. Each OTU was annotated by Mothur (v1.27) (92) with classify.seqs command, and sliva.nr_v128.align was selected as the reference database. The OTU table was resampled to the same sequence before further analysis by R 3.5.0 with the resample package.

### Statistical analysis

To compare the soil bacterial samples from different climate regions, we divided the soil samples into alpine samples and temperate samples according to collection sites. The altitude of the alpine biome sampling sites ranged from 2,796 to 4,891 m, and that of temperate biome was from 10 to 1,796 m. According to the sampling position in the soil layers, samples were divided into topsoil (0-5 cm) samples and subsoil (5-20 cm) samples. The geographic distance between sites was calculated based on geographic coordinates by the Euclidean distance method using the vegan package of R. Plant communities were classified into four functional groups (grasses, sedges, legumes, and forbs) and plant communities’ similarity and dissimilarity were calculated based on Bray-Curtis distance by the vegan package of R. Environmental factors were divided into relatively long-term environmental variables that remain relatively stable at least within a year, representing historical contingencies, and relatively short-term environmental variables that are dynamic within a year, reflecting contemporary disturbances. Relatively long-term environmental variables included MAP, MAT, pH, SOC, TN, and TP, while relatively short-term environmental variables included SWC, AP, DOC, DON, NH_4_^+^, and NO_3_^−^.

The Bray-Curtis similarity and dissimilarity of the prokaryotic community were calculated using OTU tables resampled to a minimum number of sequences from each sample (7500 in this study). The Mantel test and Partial Mantel test based on a Pearson correlation were used to test the relationship of soil prokaryotic similarity, geographic distance, and long-term multiple environmental factors or short-term multiple environmental factors. The turnover rate was estimated by the slope of the linear regression model based on the least square method. The tipping point was calculated by the function of *d*(*Y*)/*d*(*x*) = 0 in binomial function. Pearson correlation was used to test the relationship of soil prokaryotic diversity with environmental variables.

The *β*NTI was used to distinguish different ecological processes, including deterministic processes (homogenous selection and heterogeneous selection), random dispersal (homogenous dispersal, dispersal limitation), drift, and diversification (50). The |*β*NTI| > 2 means community was constructed by deterministic processes, and *β*NTI < −2 means homogenous selection plays a major role, while *β*NTI > +2 means heterogenous is more important. The −2 < *β*NTI < +2 means stochastic processes determined community succession (93). A *β*NTI analysis was performed by R 3.5.0 with the ape package. The estimation of immigration rate (*m*) was calculated by TeTame 2.0 (94) based on Hubbell’s neutral theory of biodiversity (95). Parameter estimation was rigorously performed by maximum-likelihood using the sampling formula developed by Etienne (96–99). This model is seen as a potentially useful null model in ecology; in this model, the species relative abundances in a guild are determined by two parameters, namely *θ* and *m*. The *θ* governs the appearance of a new species in the regional species pool, and *m* governs immigration into local communities of individuals from the regional species pool. We further used SEM to disentangle the causal pathways through which geographic distance, short-term environmental distance, long-term environmental distance, and plants’ community dissimilarity influence soil prokaryotic similarity. SEM in this study is implemented by AMOS software.

## Acknowledgement

This work was financially supported by the Strategic Priority Research and Program A (XDA20050104 and XDA1907304) and Program B (XDB15010201) of the Chinese Academy of Sciences, The Second Tibetan Plateau Scientific Expedition and Research (STEP) program (Grant No. 2019QZKK0304) and Chinese National Science Foundation (42041005).

## Additional information

Supplementary information is available for this paper. Reprints and permissions information is available at http://www.nature.com/reprints.

## Supplementary

**Table S1.** The mantel correlation of single environment factor with prokaryotic community similarity based on Bray-Curtis distance

|  | All |  |  |  | Alpine |  |  |  | Temperate |  |  |  | Alpine × Temperate |  |  |  |
| --- | --- | --- | --- | --- | --- | --- | --- | --- | --- | --- | --- | --- | --- | --- | --- | --- |
|  | Topsoil |  | Subsoil |  | Topsoil |  | Subsoil |  | Topsoil |  | Subsoil |  | Topsoil |  | Subsoil |  |
|  | r | p | r | p | r | p | r | p | r | p | r | p | r | p | r | p |
| MAP | 0.276 | <0.001 | 0.259 | <0.001 | 0.274 | <0.001 | 0.280 | <0.001 | 0.331 | <0.001 | -0.006 | 0.797 | 0.157 | <0.001 | 0.128 | <0.001 |
| MAT | -0.027 | 0.013 | -0.008 | 0.441 | 0.022 | 0.343 | -0.099 | <0.001 | -0.063 | 0.004 | -0.035 | 0.109 | -0.039 | 0.013 | -0.006 | 0.694 |
| pH | 0.137 | <0.001 | 0.094 | <0.001 | 0.253 | <0.001 | 0.204 | <0.001 | 0.053 | 0.016 | 0.023 | 0.289 | -0.042 | 0.006 | -0.027 | 0.084 |
| SOC | 0.274 | <0.001 | 0.191 | <0.001 | 0.420 | <0.001 | 0.277 | <0.001 | 0.124 | <0.001 | 0.067 | 0.002 | 0.187 | <0.001 | 0.148 | <0.001 |
| TN | 0.265 | <0.001 | 0.170 | <0.001 | 0.387 | <0.001 | 0.254 | <0.001 | 0.115 | <0.001 | 0.060 | 0.006 | 0.193 | <0.001 | 0.137 | <0.001 |
| TP | 0.028 | <0.001 | 0.030 | 0.006 | -0.000 | 0.985 | 0.027 | 0.240 | 0.118 | <0.001 | 0.094 | <0.001 | -0.146 | <0.001 | -0.124 | <0.001 |
| Long-term environment factors | 0.337 | 0.001 | 0.291 | 0.001 | 0.363 | 0.001 | 0.322 | 0.001 | 0.328 | 0.001 | 0.002 | 0.447 | 0.181 | 0.002 | 0.154 | 0.006 |
| SWC | 0.353 | <0.001 | 0.332 | <0.001 | 0.602 | <0.001 | 0.553 | <0.001 | 0.093 | <0.001 | 0.003 | 0.909 | 0.264 | <0.001 | 0.241 | <0.001 |
| AP | 0.185 | <0.001 | 0.074 | <0.001 | 0.355 | <0.001 | 0.145 | <0.001 | 0.002 | 0.912 | 0.080 | <0.001 | 0.197 | <0.001 | 0.080 | <0.001 |
| DOC | 0.230 | <0.001 | 0.145 | <0.001 | 0.350 | <0.001 | 0.279 | <0.001 | 0.052 | 0.017 | 0.002 | 0.913 | 0.195 | <0.001 | 0.088 | <0.001 |
| DON | 0.397 | <0.001 | 0.330 | <0.001 | 0.334 | <0.001 | 0.260 | <0.001 | 0.073 | <0.001 | 0.035 | 0.116 | 0.199 | <0.001 | 0.159 | <0.001 |
| NH<sub>4</sub><sup>+</sup> | 0.217 | <0.001 | 0.109 | <0.001 | 0.284 | <0.001 | 0.077 | <0.001 | 0.018 | 0.408 | 0.026 | 0.236 | 0.109 | <0.001 | 0.092 | <0.001 |
| NO<sub>3</sub><sup>-</sup> | 0.222 | <0.001 | 0.111 | <0.001 | 0.121 | <0.001 | 0.078 | <0.001 | 0.048 | 0.027 | 0.053 | 0.016 | 0.008 | 0.611 | -0.002 | 0.920 |
| Short-term environment factors | 0.391 | 0.001 | 0.294 | 0.001 | 0.367 | 0.001 | 0.305 | 0.001 | 0.104 | 0.091 | 0.039 | 0.251 | 0.258 | 0.001 | 0.216 | 0.001 |

**Table S2.** Partial Mantel test for correlations between prokaryotic community similarity based on Euclidean distance and each variable within long-term or short-term environmental factors in topsoil or subsoil.

|  |  | All |  |  |  | Alpine |  |  |  | Temperate |  |  |  | Alpine × Temperate |  |  |  |
| --- | --- | --- | --- | --- | --- | --- | --- | --- | --- | --- | --- | --- | --- | --- | --- | --- | --- |
|  |  | Topsoil |  | Subsoil |  | Topsoil |  | Subsoil |  | Topsoil |  | Subsoil |  | Topsoil |  | Subsoil |  |
|  |  | r | p | r | p | r | p | r | p | r | p | r | p | r | p | r | p |
|  | MAP | 0.251 | <0.001 | 0.241 | <0.001 | 0.249 | 0.002 | 0.253 | 0.003 | 0.334 | <0.001 | -0.010 | 0.546 | 0.176 | <0.001 | 0.149 | <0.001 |
|  | MAT | -0.039 | 0.984 | -0.016 | 0.740 | -0.003 | 0.518 | -0.131 | 0.991 | -0.064 | 0.983 | -0.043 | 0.876 | -0.065 | 0.980 | 0.003 | 0.439 |
|  | pH | 0.072 | 0.092 | 0.067 | 0.115 | 0.129 | 0.070 | 0.194 | 0.025 | 0.052 | 0.215 | 0.011 | 0.377 | 0.064 | 0.139 | 0.028 | 0.302 |
|  | SOC | 0.193 | <0.001 | 0.144 | 0.004 | 0.269 | <0.001 | 0.152 | 0.009 | 0.128 | 0.022 | 0.049 | 0.210 | 0.089 | 0.021 | 0.064 | 0.057 |
|  | TN | 0.197 | <0.001 | 0.131 | 0.007 | 0.239 | <0.001 | 0.150 | 0.003 | 0.117 | 0.021 | 0.042 | 0.237 | 0.116 | 0.007 | 0.101 | 0.016 |
|  | TP | 0.017 | 0.358 | 0.030 | 0.278 | -0.022 | 0.576 | 0.035 | 0.265 | 0.119 | 0.056 | 0.080 | 0.123 | -0.232 | 1.000 | -0.181 | 0.996 |
| Long-term environment factors |  | 0.290 | 0.001 | 0.262 | 0.001 | 0.306 | <0.001 | 0.272 | 0.003 | 0.329 | 0.001 | -0.017 | 0.562 | 0.203 | 0.001 | 0.159 | 0.003 |
|  | SWC | 0.265 | <0.001 | 0.266 | <0.001 | 0.534 | <0.001 | 0.488 | <0.001 | 0.008 | 0.407 | 0.002 | 0.435 | 0.167 | 0.002 | 0.225 | <0.001 |
|  | AP | 0.143 | 0.004 | 0.065 | 0.093 | 0.297 | <0.001 | 0.114 | 0.056 | -0.010 | 0.515 | 0.080 | 0.129 | 0.163 | <0.001 | 0.038 | 0.231 |
|  | DOC | 0.187 | <0.001 | 0.114 | 0.007 | 0.285 | <0.001 | 0.207 | 0.010 | 0.059 | 0.034 | 0.002 | 0.407 | 0.221 | <0.001 | 0.117 | 0.027 |
|  | DON | 0.365 | <0.001 | 0.286 | <0.001 | 0.278 | <0.001 | 0.211 | 0.002 | -0.089 | 0.919 | 0.037 | 0.247 | 0.225 | <0.001 | 0.200 | 0.002 |
|  | NH<sub>4</sub><sup>+</sup> | 0.143 | 0.006 | 0.083 | 0.078 | 0.223 | <0.001 | 0.053 | 0.241 | -0.071 | 0.908 | 0.026 | 0.315 | 0.026 | 0.331 | 0.065 | 0.093 |
|  | NO<sub>3</sub><sup>-</sup> | 0.202 | <0.001 | 0.094 | 0.032 | 0.133 | 0.031 | 0.091 | 0.073 | 0.018 | 0.368 | 0.053 | 0.224 | 0.068 | 0.140 | 0.024 | 0.323 |
| Short-term environment factors |  | 0.353 | 0.001 | 0.266 | 0.001 | 0.311 | <0.001 | 0.250 | 0.002 | -0.107 | 0.968 | 0.042 | 0.276 | 0.273 | 0.001 | 0.219 | 0.001 |

**Table S3.** Partial correlation of single environment factor with prokaryotic community similarity based on Euclidean distance at the scale of 0-1,920 km and 1,920 - 4,000 km in top-soil, and at the scale of 0-1,760 km and 1,760 - 4,000 km in sub-soil.

|  | Topsoil |  |  |  | Subsoil |  |  |  |
| --- | --- | --- | --- | --- | --- | --- | --- | --- |
|  | 0 - 1,920 |  | 1,920 - 4,000 |  | 0 - 1,760 |  | 1,760 - 4,000 |  |
|  | r | p | r | p | r | p | r | p |
| MAP | 0.194 | <0.001 | 0.177 | <0.001 | 0.207 | <0.001 | 0.130 | <0.001 |
| MAT | -0.125 | <0.001 | 0.059 | 0.001 | -0.152 | <0.001 | 0.027 | 0.119 |
| pH | 0.101 | <0.001 | 0.065 | <0.001 | 0.100 | <0.001 | 0.068 | <0.001 |
| SOC | 0.199 | <0.001 | 0.288 | <0.001 | 0.091 | <0.001 | 0.204 | <0.001 |
| TN | 0.159 | <0.001 | 0.244 | <0.001 | 0.023 | 0.114 | 0.156 | <0.001 |
| TP | 0.065 | <0.001 | 0.047 | 0.011 | 0.096 | <0.001 | 0.071 | <0.001 |
| Long-term Environment Factors | 0.272 | <0.001 | 0.274 | <0.001 | 0.268 | <0.001 | 0.223 | <0.001 |
| SWC | 0.187 | <0.001 | 0.306 | <0.001 | 0.095 | <0.001 | 0.193 | <0.001 |
| AP | 0.154 | <0.001 | 0.214 | <0.001 | 0.132 | <0.001 | 0.118 | <0.001 |
| DOC | 0.320 | <0.001 | 0.318 | <0.001 | 0.167 | <0.001 | 0.221 | <0.001 |
| DON | 0.311 | <0.001 | 0.257 | <0.001 | 0.241 | <0.001 | 0.219 | <0.001 |
| NH<sub>4</sub><sup>+</sup> | 0.269 | <0.001 | 0.219 | <0.001 | 0.065 | <0.001 | 0.065 | <0.001 |
| NO<sub>3</sub><sup>-</sup> | 0.104 | <0.001 | 0.013 | 0.479 | 0.036 | 0.013 | 0.079 | <0.001 |
| Short-term Environment Factors | 0.377 | <0.001 | 0.267 | <0.001 | 0.275 | <0.001 | 0.180 | <0.001 |

**Figure S1.**
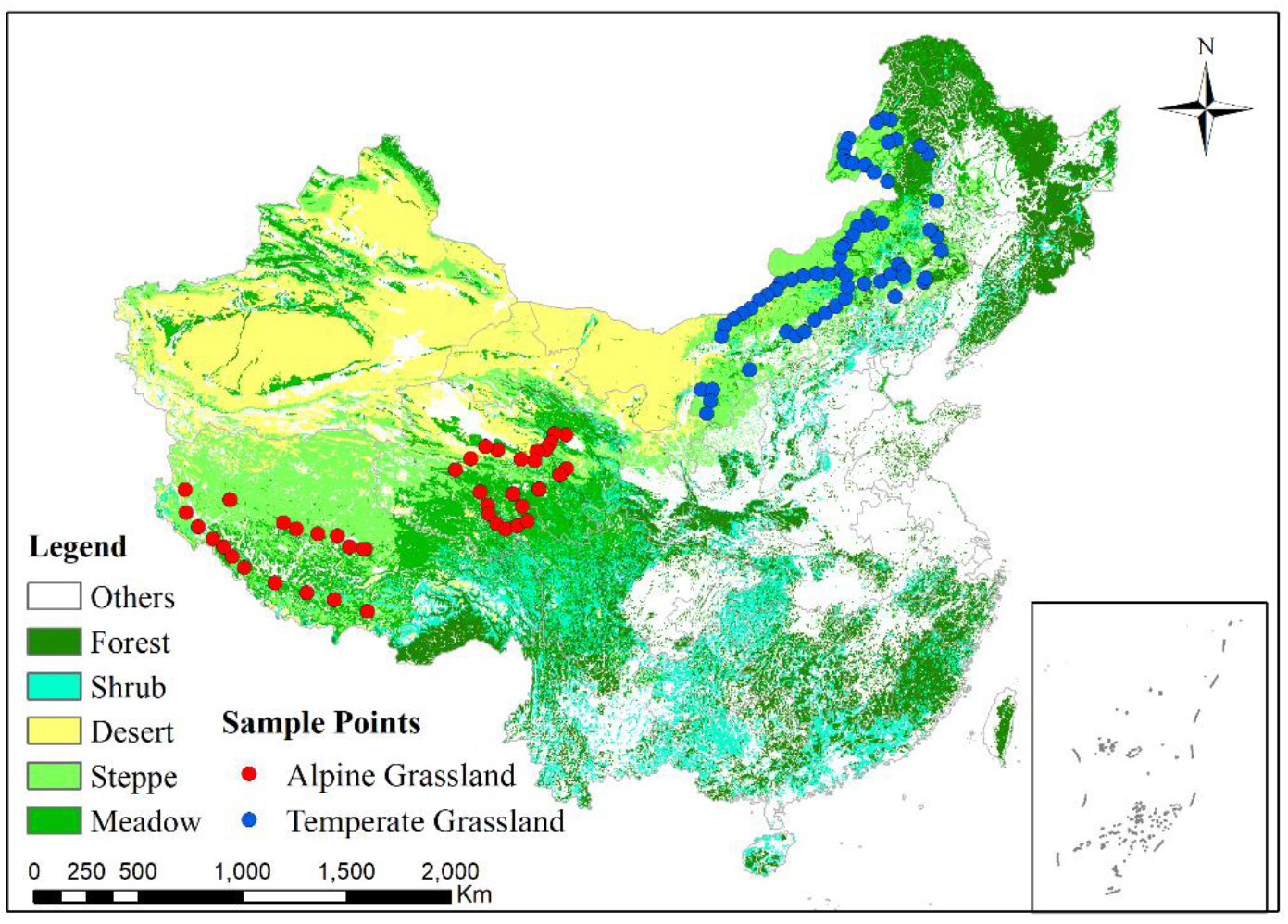
Sampling sites across 1,921 km of the alpine grassland in Qinghai-Tibet Plateau (in red) and 1,661 km of the temperate grassland in Inner Mongolia (in blue).

**Figure S2.**
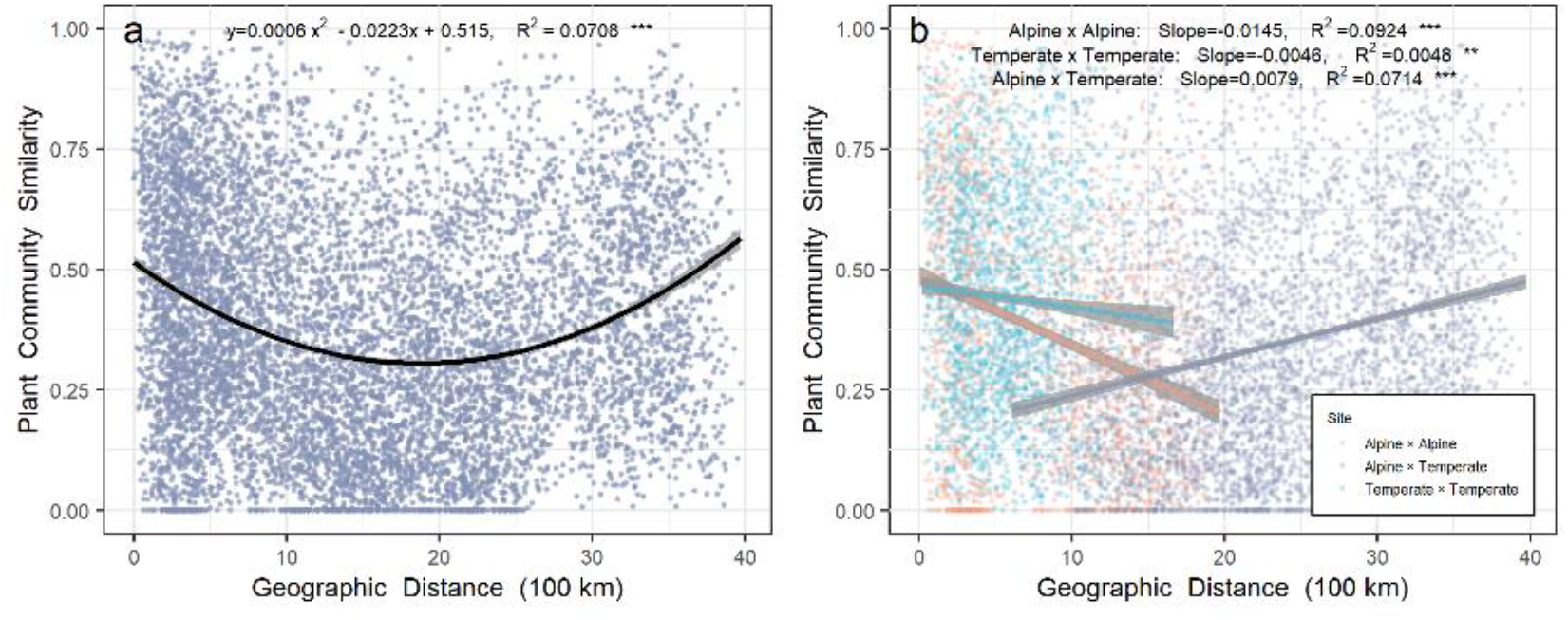
Relationship between plant community similarity over geographic distance in northern grassland of China. Light blue points are for pairwise sites in the alpine grassland. Orange points are for pairwise sites in the temperate grassland. Grey points are for pairwise sites between the alpine grassland cross temperate grassland. Grey shades stand for 95% confidence interval.

**Figure S3.**
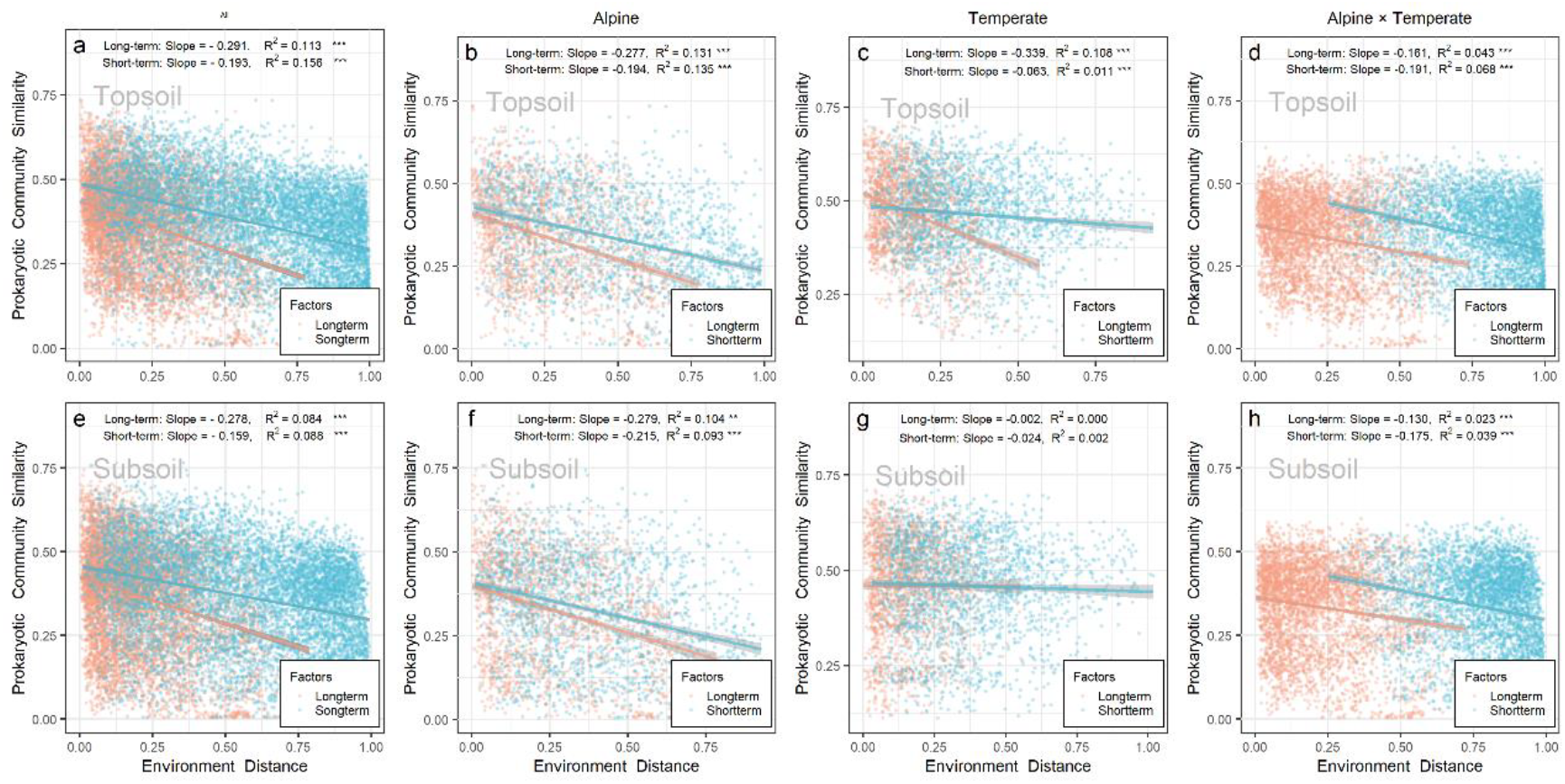
Distance decay relationship for prokaryotic community similarity over relative long-term (orange points) and short-term (light blue points) environmental factors. (a-d) topsoil; (e-h) subsoil. Shades stand for 95% confidence interval.

**Figure S4.**
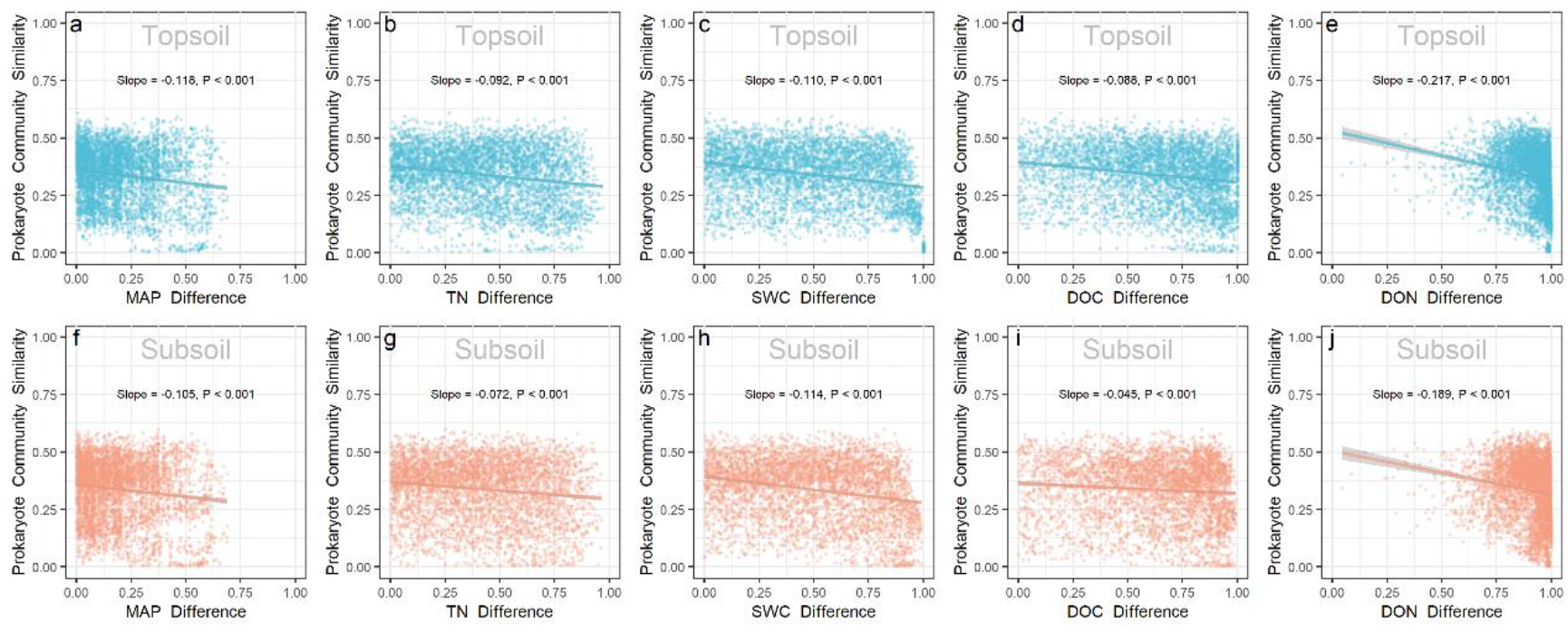
The scatter plots of the mantel correlation for prokaryotic community similarity with individual environment factor (MAP, TN, SWC, DOC, and DON) Bray-Curtis distance. Panels **a-e** were for topsoil, and panel **f-j** for subsoil.

**Figure S5.**
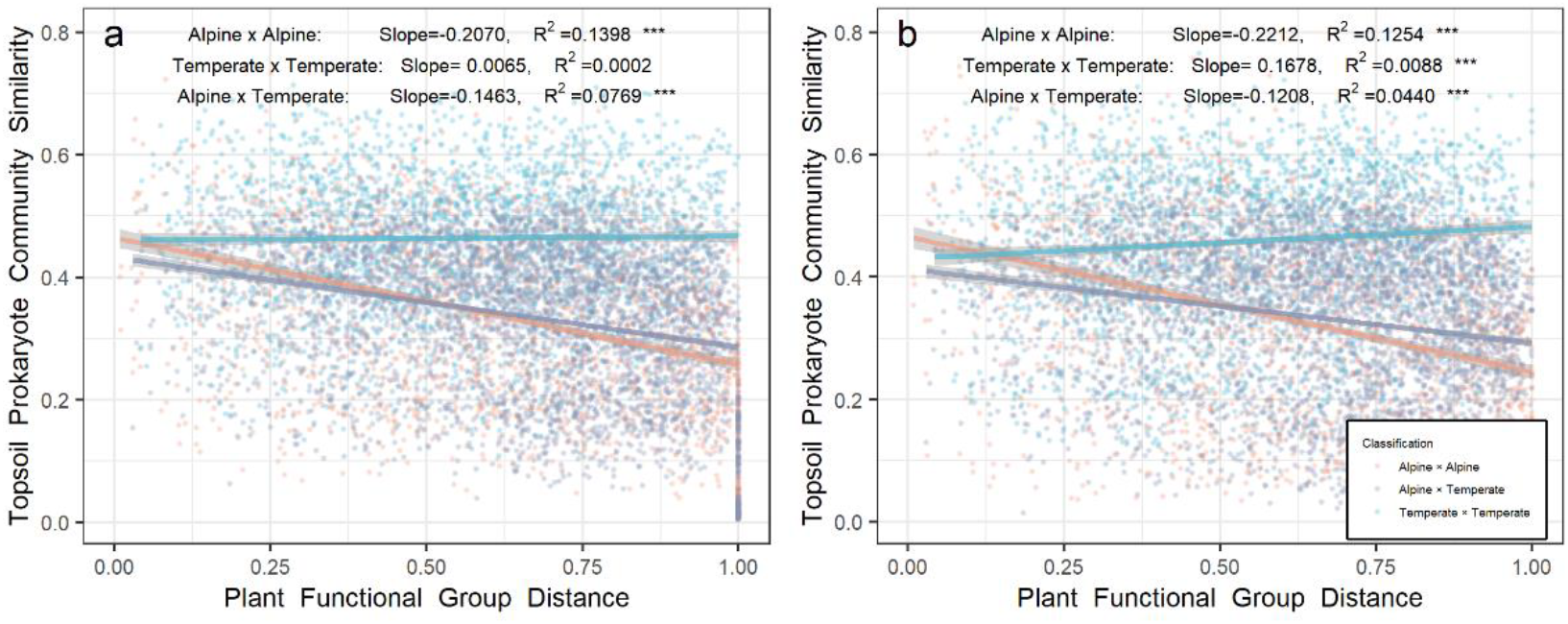
Relationship between prokaryote community over plant community dissimilarity. (a) topsoil; (b)subsoil. Light blue points are for pairwise sites in the alpine grassland. Orange points are for pairwise sites in the temperate grassland. Grey points are for pairwise sites between the alpine grassland cross temperate grassland. Grey shades stand for 95% confidence interval.

**Figure S6.**
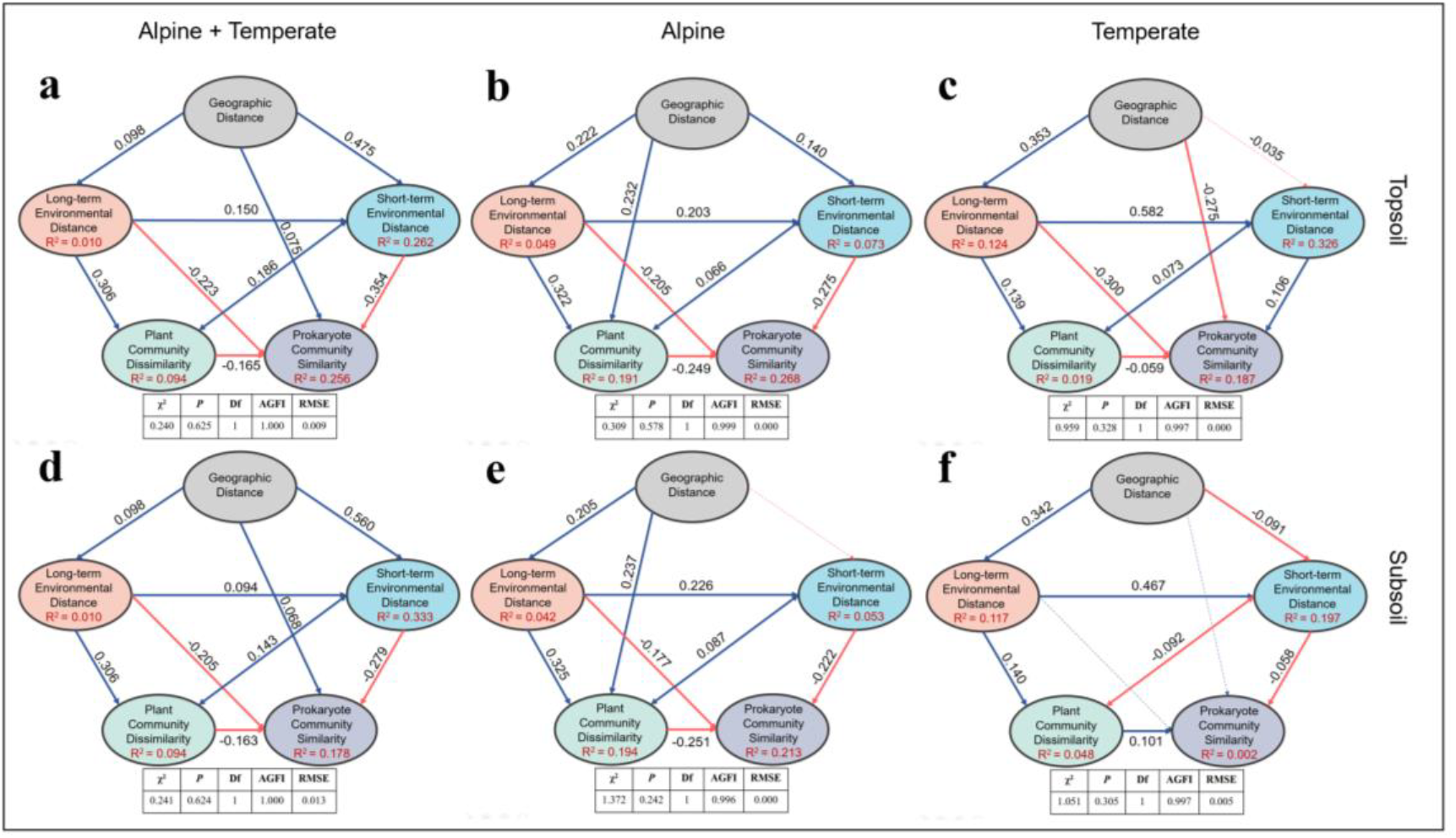
Structure equation model for geographic distance (based on Euclidean index), long-term environmental distance, short-term environmental distance, and plant community dissimilarity based on Bray-Curtis distance in affecting soil prokaryote community similarity. **(a)** topsoil of the northern grassland; **(b)** topsoil of the alpine grassland; **(c)** topsoil of the temperate grassland; **(d)** subsoil of the northern grassland; **(e)** subsoil of the alpine grassland; **(f)** subsoil of the temperate grassland. Red lines stand for negative correlation and blue lines stand for positive correlation; Bold lines stand for positive correlation; Bold lines stand for significance at 0.05 level.

**Figure S7.**
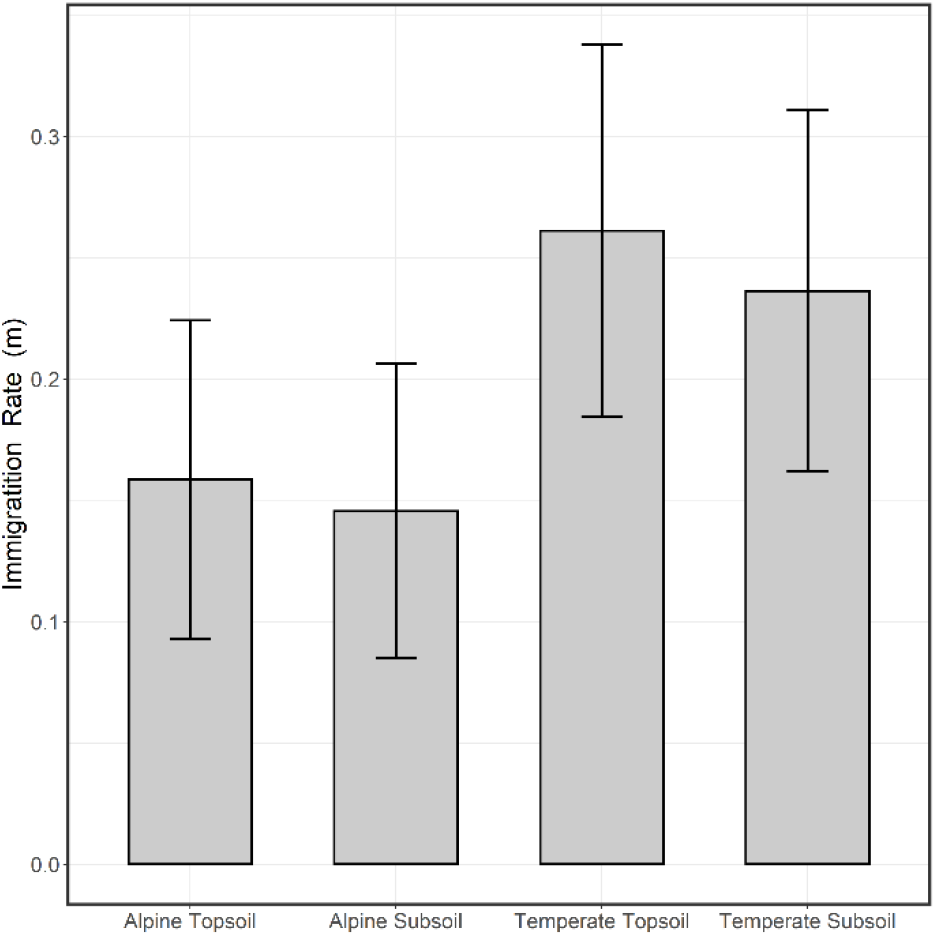
The immigration rate (m) of soil prokaryote community in topsoil and subsoil of the alpine and temperate grassland biomes.

## References

1. Bahram M, Koljalg U, Courty P-E, Diedhiou AG, Kjoller R, Polme S, et al. The distance decay of similarity in communities of ectomycorrhizal fungi in different ecosystems and scales. J Ecol. 2013;101(5):1335–44.

2. Hanson CA, Fuhrman JA, Horner-Devine MC, Martiny JBH. Beyond biogeographic patterns: processes shaping the microbial landscape. Nat Rev Microbiol. 2012;10(7):497–506.

3. Condit R, Pitman N, Leigh EG, Chave J, Terborgh J, Foster RB, et al. Beta-diversity in tropical forest trees. Science. 2002;295(5555):666–9.

4. Tuomisto H, Ruokolainen K, Yli-Halla M. Dispersal, environment, and floristic variation of western Amazonian forests. Science. 2003;299(5604):241–4.

5. Gilbert B, Lechowicz MJ. Neutrality, niches, and dispersal in a temperate forest understory. Proceedings of the National Academy of Sciences of the United States of America. 2004;101(20):7651–6.

6. Dexter KG, Terborgh JW, Cunningham CW. Historical effects on beta diversity and community assembly in Amazonian trees. Proceedings of the National Academy of Sciences of the United States of America. 2012;109(20):7787–92.

7. Zhao Z, Li S, Liu J, Peng J, Wang Y. The distance decay of similarity in climate variation and vegetation dynamics. Environ Earth Sci. 2015;73(8):4659–70.

8. Draper FC, Baraloto C, Brodrick PG, Phillips OL, Vasquez Martinez R, Honorio Coronado EN, et al. Imaging spectroscopy predicts variable distance decay across contrasting Amazonian tree communities. J Ecol. 2019;107(2):696–710.

9. Gomez-Rodriguez C, Baselga A. Variation among European beetle taxa in patterns of distance decay of similarity suggests a major role of dispersal processes. Ecography. 2018;41(11):1825–34.

10. van der Mescht L, Warburton EM, Khokhlova IS, Stanko M, Vinarski MV, Korallo-Vinarskaya NP, et al. Biogeography of parasite abundance: latitudinal gradient and distance decay of similarity in the abundance of fleas and mites, parasitic on small mammals in the Palearctic, at three spatial scales. Int J Parasitol. 2018;48(11):857–66.

11. Warfe DM, Pettit NE, Magierowski RH, Pusey BJ, Davies PM, Douglas MM, et al. Hydrological connectivity structures concordant plant and animal assemblages according to niche rather than dispersal processes. Freshwater Biol. 2013;58(2):292–305.

12. Beisner BE, Peres PR, Lindstrom ES, Barnett A, Longhi ML. The role of environmental and spatial processes in structuring lake communities from bacteria to fish. Ecology. 2006;87(12):2985–91.

13. Matsuoka S, Kawaguchi E, Osono T. Temporal distance decay of similarity of ectomycorrhizal fungal community composition in a subtropical evergreen forest in Japan. Fems Microbiol Ecol. 2016;92(5).

14. Oono R, Rasmussen A, Lefevre E. Distance decay relationships in foliar fungal endophytes are driven by rare taxa. Environ Microbiol. 2017;19(7):2794–805.

15. Wang XB, Lu XT, Yao J, Wang ZW, Deng Y, Cheng WX, et al. Habitat-specific patterns and drivers of bacterial beta-diversity in China’s drylands. Isme J. 2017;11(6):1345–58.

16. Finkel OM, Burch AY, Elad T, Huse SM, Lindow SE, Post AF, et al. Distance-Decay Relationships Partially Determine Diversity Patterns of Phyllosphere Bacteria on Tamrix Trees across the Sonoran Desert. Appl Environ Microb. 2012;78(17):6187–93.

17. Jiao S, Yang Y, Xu Y, Zhang J, Lu Y. Balance between community assembly processes mediates species coexistence in agricultural soil microbiomes across eastern China. Isme J. 2020;14(1):202–16.

18. Jiao S, Xu Y, Zhang J, Lu Y. Environmental filtering drives distinct continental atlases of soil archaea between dryland and wetland agricultural ecosystems. Microbiome. 2019;7.

19. Shi Y, Li YT, Xiang XJ, Sun RB, Yang T, He D, et al. Spatial scale affects the relative role of stochasticity versus determinism in soil bacterial communities in wheat fields across the North China Plain. Microbiome. 2018;6.

20. Zinger L, Boetius A, Ramette A. Bacterial taxa-area and distance-decay relationships in marine environments. Mol Ecol. 2014;23(4):954–64.

21. Barreto DP, Conrad R, Klose M, Claus P, Enrich-Prast A. Distance-Decay and Taxa-Area Relationships for Bacteria, Archaea and Methanogenic Archaea in a Tropical Lake Sediment. Plos One. 2014;9(10).

22. Shi Y, Adams JM, Ni Y, Yang T, Jing X, Chen L, et al. The biogeography of soil archaeal communities on the eastern Tibetan Plateau. Sci Rep-Uk. 2016;6.

23. Chen W, Ren K, Isabwe A, Chen H, Liu M, Yang J. Stochastic processes shape microeukaryotic community assembly in a subtropical river across wet and dry seasons. Microbiome. 2019;7(1).

24. Wang J, Zhang T, Li L, Li J, Feng Y, Lu Q. The Patterns and Drivers of Bacterial and Fungal beta-Diversity in a Typical Dryland Ecosystem of Northwest China. Front Microbiol. 2017;8.

25. Yang T, Adams JM, Shi Y, Sun H, Cheng L, Zhang Y, et al. Fungal community assemblages in a high elevation desert environment: Absence of dispersal limitation and edaphic effects in surface soil. Soil Biol Biochem. 2017;115:393–402.

26. Zhang J, Zhang B, Liu Y, Guo Y, Shi P, Wei G. Distinct large-scale biogeographic patterns of fungal communities in bulk soil and soybean rhizosphere in China. Sci Total Environ. 2018;644:791–800.

27. Hu H-W, Zhang L-M, Yuan C-L, Zheng Y, Wang J-T, Chen D, et al. The large-scale distribution of ammonia oxidizers in paddy soils is driven by soil pH, geographic distance, and climatic factors. Front Microbiol. 2015;6.

28. Angermeyer A, Crosby SC, Huber JA. Decoupled distance-decay patterns between dsrA and 16S rRNA genes among salt marsh sulfate-reducing bacteria. Environ Microbiol. 2016;18(1):75–86.

29. Zhou JZ, Kang S, Schadt CW, Garten CT. Spatial scaling of functional gene diversity across various microbial taxa. Proceedings of the National Academy of Sciences of the United States of America. 2008;105(22):7768–73.

30. Zhou J, Deng Y, Shen L, Wen C, Yan Q, Ning D, et al. Temperature mediates continental-scale diversity of microbes in forest soils. Nat Commun. 2016;7.

31. Ribeiro KF, Duarte L, Crossetti LO. Everything is not everywhere: a tale on the biogeography of cyanobacteria. Hydrobiologia. 2018;820(1):23–48.

32. Soininen J, McDonald R, Hillebrand H. The distance decay of similarity in ecological communities. Ecography. 2007;30(1):3–12.

33. Cottenie K. Integrating environmental and spatial processes in ecological community dynamics. Ecology Letters. 2005;8(11):1175–82.

34. Bell T. Experimental tests of the bacterial distance-decay relationship. Isme Journal. 2010;4(11):1357–65.

35. Lindstrom ES, Ostman O. The Importance of Dispersal for Bacterial Community Composition and Functioning. Plos One. 2011;6(10).

36. Leibold MA, Holyoak M, Mouquet N, Amarasekare P, Chase JM, Hoopes MF, et al. The metacommunity concept: a framework for multi-scale community ecology. Ecology Letters. 2004;7(7):601–13.

37. Stegen JC, Lin X, Konopka AE, Fredrickson JK. Stochastic and deterministic assembly processes in subsurface microbial communities. Isme J. 2012;6(9):1653–64.

38. Chase JM, Myers JA. Disentangling the importance of ecological niches from stochastic processes across scales. Philos T R Soc B. 2011;366(1576):2351–63.

39. Hubbell SP. Neutral theory in community ecology and the hypothesis of functional equivalence. Functional Ecology. 2005;19(1):166–72.

40. Feng M, Tripathi BM, Shi Y, Adams JM, Zhu Y-G, Chu H. Interpreting distance-decay pattern of soil bacteria via quantifying the assembly processes at multiple spatial scales. Microbiologyopen. 2019;8(9).

41. Chu H, Sun H, Tripathi BM, Adams JM, Huang R, Zhang Y, et al. Bacterial community dissimilarity between the surface and subsurface soils equals horizontal differences over several kilometers in the western Tibetan Plateau. Environ Microbiol. 2016;18(5):1523–33.

42. Green JL, Holmes AJ, Westoby M, Oliver I, Briscoe D, Dangerfield M, et al. Spatial scaling of microbial eukaryote diversity. Nature. 2004;432(7018):747–50.

43. Whitaker RJ, Grogan DW, Taylor JW. Geographic barriers isolate endemic populations of hyperthermophilic archaea. Science. 2003;301(5635):976–8.

44. Okie JG, Van Horn DJ, Storch D, Barrett JE, Gooseff MN, Kopsova L, et al. Niche and metabolic principles explain patterns of diversity and distribution: theory and a case study with soil bacterial communities. Proceedings of the Royal Society B-Biological Sciences. 2015;282(1809).

45. Tobler WR, Mielke HW, Detwyler TR. Geobotanical distance between New-Zealand and neighboring island. Bioscience. 1970;20(9):537–&.

46. Nekola JC, White PS. The distance decay of similarity in biogeography and ecology. Journal of Biogeography. 1999;26(4):867–78.

47. Wang X, Wen X, Deng Y, Xia Y, Yang Y, Zhou J. Distance-Decay Relationship for Biological Wastewater Treatment Plants. Appl Environ Microb. 2016;82(16):4860–6.

48. Leff JW, Bardgett RD, Wilkinson A, Jackson BG, Pritchard WJ, De Long JR, et al. Predicting the structure of soil communities from plant community taxonomy, phylogeny, and traits. Isme J. 2018;12(7):1794–805.

49. Xing R, Gao Q-b, Zhang F-q, Wang J-l, Chen S-l. Large-scale distribution of bacterial communities in the Qaidam Basin of the Qinghai-Tibet Plateau. Microbiologyopen. 2019;8(10).

50. Zhou J, Ning D. Stochastic Community Assembly: Does It Matter in Microbial Ecology? Microbiology and Molecular Biology Reviews. 2017;81(4).

51. Zhang B, Xue K, Zhou S, Che R, Du J, Tang L, et al. Phosphorus mediates soil prokaryote distribution pattern along a small-scale elevation gradient in Noijin Kangsang Peak, Tibetan Plateau. Fems Microbiol Ecol. 2019;95(6).

52. Suseela V, Conant RT, Wallenstein MD, Dukes JS. Effects of soil moisture on the temperature sensitivity of heterotrophic respiration vary seasonally in an old-field climate change experiment. Global Change Biology. 2012;18(1):336–48.

53. Moyano FE, Manzoni S, Chenu C. Responses of soil heterotrophic respiration to moisture availability: An exploration of processes and models. Soil Biol Biochem. 2013;59:72–85.

54. Kaye JP, Hart SC. Competition for nitrogen between plants and soil microorganisms. Trends in Ecology & Evolution. 1997;12(4):139–43.

55. Brockett BFT, Prescott CE, Grayston SJ. Soil moisture is the major factor influencing microbial community structure and enzyme activities across seven biogeoclimatic zones in western Canada. Soil Biol Biochem. 2012;44(1):9–20.

56. Mansson KF, Olsson MO, Falkengren-Grerup U, Bengtsson G. Soil moisture variations affect short-term plant-microbial competition for ammonium, glycine, and glutamate. Ecol Evol. 2014;4(7):1061–72.

57. Li Y, Adams J, Shi Y, Wang H, He J-S, Chu H. Distinct Soil Microbial Communities in habitats of differing soil water balance on the Tibetan Plateau. Sci Rep-Uk. 2017;7.

58. Wichern F, Joergensen RG. Soil Microbial Properties Along a Precipitation Transect in Southern Africa. Arid Land Research and Management. 2009;23(2):115–26.

59. Hu Y, Xiang D, Veresoglou SD, Chen F, Chen Y, Hao Z, et al. Soil organic carbon and soil structure are driving microbial abundance and community composition across the arid and semi-arid grasslands in northern China. Soil Biol Biochem. 2014;77:51–7.

60. Chen D, Mi J, Chu P, Cheng J, Zhang L, Pan Q, et al. Patterns and drivers of soil microbial communities along a precipitation gradient on the Mongolian Plateau. Landscape Ecol. 2015;30(9):1669–82.

61. Zeng J, Liu X, Song L, Lin X, Zhang H, Shen C, et al. Nitrogen fertilization directly affects soil bacterial diversity and indirectly affects bacterial community composition. Soil Biol Biochem. 2016;92:41–9.

62. He D, Xiang X, He J-S, Wang C, Cao G, Adams J, et al. Composition of the soil fungal community is more sensitive to phosphorus than nitrogen addition in the alpine meadow on the Qinghai-Tibetan Plateau. Biol Fert Soils. 2016;52(8):1059–72.

63. Alden L, Demoling F, Baath E. Rapid method of determining factors limiting bacterial growth in soil. Appl Environ Microb. 2001;67(4):1830–8.

64. Drenovsky RE, Vo D, Graham KJ, Scow KM. Soil water content and organic carbon availability are major determinants of soil microbial community composition. Microbial Ecology. 2004;48(3):424–30.

65. de Vries FT, Manning P, Tallowin JRB, Mortimer SR, Pilgrim ES, Harrison KA, et al. Abiotic drivers and plant traits explain landscape-scale patterns in soil microbial communities. Ecology Letters. 2012;15(11):1230–9.

66. Grayston SJ, Wang SQ, Campbell CD, Edwards AC. Selective influence of plant species on microbial diversity in the rhizosphere. Soil Biol Biochem. 1998;30(3):369–78.

67. Singh BK, Millard P, Whiteley AS, Murrell JC. Unravelling rhizosphere-microbial interactions: opportunities and limitations. Trends Microbiol. 2004;12(8):386–93.

68. Berg G, Smalla K. Plant species and soil type cooperatively shape the structure and function of microbial communities in the rhizosphere. Fems Microbiol Ecol. 2009;68(1):1–13.

69. Hiiesalu I, Paertel M, Davison J, Gerhold P, Metsis M, Moora M, et al. Species richness of arbuscular mycorrhizal fungi: associations with grassland plant richness and biomass. New Phytologist. 2014;203(1):233–44.

70. Peay KG, Baraloto C, Fine PVA. Strong coupling of plant and fungal community structure across western Amazonian rainforests. Isme J. 2013;7(9):1852–61.

71. Prober SM, Leff JW, Bates ST, Borer ET, Firn J, Harpole WS, et al. Plant diversity predicts beta but not alpha diversity of soil microbes across grasslands worldwide. Ecology Letters. 2015;18(1):85–95.

72. van der Heijen MGA. The unseen majority: Soil microbes as drivers of plant diversity and productivity in terrestrial ecosystems (vol 11, pg 296, 2008). Ecology Letters. 2008;11(6):651-.

73. Badri DV, Vivanco JM. Regulation and function of root exudates. Plant Cell Environ. 2009;32(6):666–81.

74. Bais HP, Weir TL, Perry LG, Gilroy S, Vivanco JM. The role of root exudates in rhizosphere interations with plants and other organisms. Annual Review of Plant Biology. 2006;57:233–66.

75. Baetz U, Martinoia E. Root exudates: the hidden part of plant defense. Trends in Plant Science. 2014;19(2):90–8.

76. Bulgarelli D, Schlaeppi K, Spaepen S, van Themaat EVL, Schulze-Lefert P. Structure and Functions of the Bacterial Microbiota of Plants. In: Merchant SS, editor. Annual Review of Plant Biology, Vol 64. Annual Review of Plant Biology. 642013. p. 807–38.

77. Hooper DU, Bignell DE, Brown VK, Brussaard L, Dangerfield JM, Wall DH, et al. Interactions between aboveground and belowground biodiversity in terrestrial ecosystems: Patterns, mechanisms, and feedbacks. Bioscience. 2000;50(12):1049–61.

78. Wardle DA. The influence of biotic interactions on soil biodiversity. Ecology Letters. 2006;9(7):870–86.

79. Mellado-Vazquez PG, Lange M, Bachmann D, Gockele A, Karlowsky S, Milcu A, et al. Plant diversity generates enhanced soil microbial access to recently photosynthesized carbon in the rhizosphere. Soil Biol Biochem. 2016;94:122–32.

80. Jones DL. Organic acids in the rhizosphere - a critical review. Plant and Soil. 1998;205(1):25–44.

81. Zhalnina K, Louie KB, Hao Z, Mansoori N, da Rocha UN, Shi S, et al. Dynamic root exudate chemistry and microbial substrate preferences drive patterns in rhizosphere microbial community assembly. Nature Microbiology. 2018;3(4):470–80.

82. Delgado-Baquerizo M, Maestre FT, Reich PB, Jeffries TC, Gaitan JJ, Encinar D, et al. Microbial diversity drives multifunctionality in terrestrial ecosystems. Nat Commun. 2016;7.

83. Xue K, Zhang B, Zhou S, Ran Q, Tang L, Che R, et al. Soil microbial communities in alpine grasslands on the Tibetan Plateau and their influencing factors. Chinese Science Bulletin. 2019;64(27):2915–27.

84. Horner-Devine MC, Lage M, Hughes JB, Bohannan BJM. A taxa-area relationship for bacteria. Nature. 2004;432(7018):750–3.

85. Martiny JBH, Eisen JA, Penn K, Allison SD, Horner-Devine MC. Drivers of bacterial beta-diversity depend on spatial scale. Proceedings of the National Academy of Sciences of the United States of America. 2011;108(19):7850–4.

86. Chase JM. Drought mediates the importance of stochastic community assembly. Proceedings of the National Academy of Sciences of the United States of America. 2007;104(44):17430–4.

87. Evans S, Martiny JBH, Allison SD. Effects of dispersal and selection on stochastic assembly in microbial communities. Isme J. 2017;11(1):176–85.

88. Louca S, Jacques SMS, Pires APF, Leal JS, Srivastava DS, Parfrey LW, et al. High taxonomic variability despite stable functional structure across microbial communities. Nature Ecology & Evolution. 2017;1(1).

89. Young JW, Locke JCW, Elowitz MB. Rate of environmental change determines stress response specificity. Proceedings of the National Academy of Sciences of the United States of America. 2013;110(10):4140–5.

90. Cox F, Newsham KK, Bol R, Dungait JAJ, Robinson CH. Not poles apart: Antarctic soil fungal communities show similarities to those of the distant Arctic. Ecology Letters. 2016;19(5):528–36.

91. Davison J, Moora M, Oepik M, Adholeya A, Ainsaar L, Ba A, et al. Global assessment of arbuscular mycorrhizal fungus diversity reveals very low endemism. Science. 2015;349(6251):970–3.

92. Schloss PD, Westcott SL, Ryabin T, Hall JR, Hartmann M, Hollister EB, et al. Introducing mothur: Open-Source, Platform-Independent, Community-Supported Software for Describing and Comparing Microbial Communities. Appl Environ Microb. 2009;75(23):7537–41.

93. Dini-Andreote F, Stegen JC, van Elsas JD, Salles JF. Disentangling mechanisms that mediate the balance between stochastic and deterministic processes in microbial succession. Proceedings of the National Academy of Sciences of the United States of America. 2015;112(11):E1326–E32.

94. Chave J, Jabot F. Estimation of neutral parameters by maximum likelihood. Université Paul Sabatier. 2008.

95. SP. H. The unified neutral theory of biodiversity and biogeography. Princeton University Press. 2001;Princeton(NJ).

96. Etienne RS. A new sampling formula for neutral biodiversity. Ecology Letters. 2005;8(3):253–60.

97. Etienne RS, Alonso D. A dispersal-limited sampling theory for species and alleles. Ecology Letters. 2005;8(11):1147–56.

98. Etienne RS, Olff H. Confronting different models of community structure to species-abundance data: a Bayesian model comparison. Ecology Letters. 2005;8(5):493–504.

99. Etienne RS, Latimer AM, Silander JA, Cowling RM. Comment on “Neutral ecological theory reveals isolation and rapid speciation in a biodiversity hot spot”. Science. 2006;311(5761):610B–+.

